# Shift from visceral to subcutaneous fat in *Cyp17a1* knockout rats prevent the progression of metabolic syndrome

**DOI:** 10.1101/2024.09.22.614373

**Authors:** Beom-Jin Jeon, Jeong-Hwa Lee, Dong-Hyeok Kwon, Hee-Kyoung Kim, Goo Jang

**Affiliations:** Laboratory of Theriogenology and Biotechnology, Department of Veterinary Clinical Science, College of Veterinary Medicine, BK21 FOUR Future Veterinary Medicine Leading Education, and the Research Institute of Veterinary Science, Seoul National University, Seoul, Republic of Korea; Comparative medicine Disease Research Center, Seoul National University, Seoul, Republic of Korea; K-BIO KIURI Center, Seoul National University, Seoul, Republic of Korea; Department of Biomedical Sciences, School of Veterinary Medicine, University of Pennsylvania, Philadelphia, PA 19104, USA; LARTBio Incorp., Gyeonggi-Do, Republic of Korea

**Author notes:** Correspondence: Goo Jang, D.V.M., Ph.D., 1 Gwanak-ro, Gwanak-gu, Seoul, Republic of Korea, 08826. These authors contributed equally (Beom-Jin Jeon, Jeong-Hwa Lee).

**Keywords:** Adipose tissue, *Cyp17a1*, Gene edited rats, Metabolic syndrome, Obesity

## Abstract

In this study, we investigated the effects of *Cyp17a1* gene knockout (KO) on obesity and metabolic syndrome. *Cyp17a1* KO in rats using CRISPR-Cas9 resulted in sex dimorphism and obesity, and interestingly, idiopathic accumulation was found in subcutaneous adipose tissue. Surprisingly, an insulin tolerance test and oral glucose tolerance test did not show any issues with insulin sensitivity and secretion despite hyperglycemia. In addition, *Cyp17a1* KO rats showed normal plasma insulin and free fatty acid levels compared to wild-type rats, and blood biochemistry analysis revealed normal triglyceride, total cholesterol, high-density lipoprotein, and low-density lipoprotein levels. *Cyp17a1* KO adipose-tissue-derived stem cells from subcutaneous fat showed increased expression of KLF5, an early adipogenesis marker, which implies enhanced adipogenic potential in subcutaneous adipose tissue. When gene expression associated with lipid, glucose, and insulin metabolism as well as inflammation in adipose tissue was examined, a metabolic shift to subcutaneous adipose tissue was discovered in the *Cyp17a1* KO group. In conclusion, in the Cyp17a1 KO rat models we generated for the first time, the phenotype promoted by obesity reflected health obesity hypothesis, but this did not result in metabolic syndrome due to enhanced metabolism in the subcutaneous fats.

## Introduction

Obesity has been a recent global health concern, leading to an increase in diabetes, high blood pressure, and cardiovascular disease, among other disorders (1). Metabolic syndrome is the term for the complex physical condition that includes diabetes and cardiovascular illnesses brought on by obesity (2–4). As the prevalence of obesity increases, there is a corresponding rise in the incidence of metabolic syndrome (5). This trend underscores the urgency of advancing research into the mechanistic pathways that link obesity to subsequent metabolic disorders such as diabetes and cardiovascular diseases (6, 7). In this respect, adipose tissue is one of the main focuses of research regarding the progression of metabolic syndrome (8, 9).

Adipose tissue is now understood to engage in cross-talk with other organs as well as within itself, functioning as a type of endocrine gland that regulates blood glucose and free fatty acid concentrations through lipolysis and lipogenesis (10). Additionally, adipose tissue secretes adipokines such as leptin and adiponectin, which are known to regulate not only energy storage but also obesity and insulin resistance (11–14). Recent studies on the role of adipose tissue in metabolic syndrome development have focused on the different roles of visceral fat and subcutaneous fat. Visceral fat is located inside the abdominal cavity and dysfunction in visceral fat can lead to metabolic diseases due to inflammation and disturbed metabolic balance (15–17). In contrast, subcutaneous fat appears to have a protective role, as its release of adipokines can prevent the progression of metabolic syndrome and prevent excess fat storage in visceral fat and ectopic depots including the liver (16–20). Considering these distinct roles of different adipose tissues, targeted research on each specific type of fat is crucial for a comprehensive understanding of metabolic syndrome and obesity (19, 21). Furthermore, considering that obesity is a major risk factor for numerous metabolic derangements and related diseases which can be caused by monogenic or polygenic factors (22), there is a compelling need to delve into the genetic architecture of obesity to better understand and potentially mitigate its impact (23).

The *Cyp17a1* gene is involved in steroidogenesis and is required for the synthesis of glucocorticoids and sex hormones (24). The *Cyp17a1* gene has recently been linked to both sex dimorphism and atherosclerosis (25–28). Furthermore, some research findings indicate that obesity develops from a mutation in the *Cyp17a1* gene (29, 30); however, the exact mechanism underlying obesity and the impact of obesity on metabolic syndrome resulting from *Cyp17a1* gene mutation remains unknown. Furthermore, not enough is known about the impact of *Cyp17a1* gene knockout (KO)-induced modifications in steroidogenesis and its effects on adipokine production and lipid metabolism in adipose tissue depot specifically.

Therefore, the goal of this research was to find out how the *Cyp17a1* gene alters adipose tissue metabolism and how this influences the development of obesity and the resulting metabolic syndrome. To achieve this, we produced *Cyp17a1* KO rats and verified their obesity phenotype, assessing key components of metabolic syndrome including blood glucose levels, blood pressure, glucose tolerance, and insulin resistance. We also separately isolated subcutaneous and visceral (perigonadal) fat to examine alterations in the metabolism of adipose tissue and adipose-derived stem cells (ADSCs).

## Materials & Methods

### Animals

All animal care and procedures were approved by the Institutional Animal Care and Use Committee (No.201222-4-2) of Seoul National University Institute of Laboratory Animal Resources and performed under the guidelines of Seoul National University. SD rats used in this study were purchased from Orient-Bio (Seoungnam, Republic of Korea) and maintained at 24±2 ℃, 50% humidity, and a 12:12 h light-dark cycle (lights on from 07:00 to 19:00).

### Superovulation and Embryo Collection

Female rats were induced to superovulate by an intraperitoneal injection of 150 IU/kg Pregnant mare serum gonadotrophin (Daesung Microbiological Labs, Republic of Korea) and 150 IU/kg human chorionic gonadotrophin (Daesung Microbiological labs) with a 48-h period, and then mated with males. Rats were anesthetized with an intramuscular injection of a 1.5 mg xylazine (Rompun®, Elanco Korea, Republic of Korea) and 0.35 mg alfaxalone (Alfaxan® multidose, Jurox, Australia) mixture and euthanized by cervical dislocation. One-cell-stage embryos were collected in M2 medium (Sigma-Aldrich, USA) from oviducts of females the day after mating, and cultured in mR1ECM medium (Cosmobio, Japan).

### gRNA Synthesis, Embryo Electroporation and Embryo Transfer

Guide RNA (gRNA) for *Cyp17a1* knockout was designed using an online tool (CRISPR RGEN tool; http://www.rgenome.net/) and synthesized using a precision gRNA synthesis kit (Invitrogen, USA). The gRNA sequences used in this study are described in S1 Fig a. A total of 200 ng/μL gRNA and Cas9 RNP (1:1) were incubated for 5 min and transfected into one-cell stage rat embryos using Genome Editor (GEB15, BEX, Japan) in conditions of 30 V, 7 times for 3 msec, and 97 msec intervals. Two-cell stage embryos were transferred to the oviducts of pseudo-pregnant recipient females.

### Embryo Cryopreservation

Embryo cryopreservation was conducted as previously described (31). Briefly, two-cell stage rat embryos were washed 3 times using M2 medium and BoviFreeze (Minitube, Germany) subsequently. Washed embryos were loaded in a Ministraw (Minitube) and frozen using Freeze Control embryo freezers (CL8800i, Minitube).

### Genotyping

Genomic DNA for genotyping was extracted from the rat tails using a DNeasy Blood & Tissue Kit (Qiagen, Germany). PCR for *Cyp17a1* amplification was conducted using Mastercycler X50a (Eppendorf, Germany). Primers used in PCR are listed in S1 Table.

### qPCR Analysis

Total RNAs were extracted from rat gonad tissues and adipose tissues using an RNeasy Mini Kit (Qiagen) and 5 µg of total RNA was used for synthesizing complementary DNA (cDNA) using an RNA to cDNA EcoDry™ Premix Kit (Clontech, USA). Quantitative PCR (qPCR) was conducted in a MicroAmp™ Optical 96-Well Reaction Plate (Applied Biosystems, USA) using TB Green^®^ Premix Ex Taq™ (TAKARA, Japan). Quantitative PCR was conducted using a Quantstudio 1 Real-Time PCR instrument (Applied Biosystems). Target-gene-related expression was normalized to *Actb* expression using the comparative CT (2^-ΔΔCt^) method. Primers used in qPCR are listed in S1 Table.

### Estradiol and Testosterone Assay

Serum estradiol and testosterone were assessed with an electrochemiluminescence immunoassay (ECLIA) Elecsys Estradiol III and Testosterone II (Roche, Switzerland) conducted by the Global Clinical Central Lab (GCCL, Republic of Korea) using a Cobas 8000 e801 (Roche). For assessment of estradiol, two biotinylated monoclonal anti-estradiol antibodies (rabbit) were used at concentrations of 2.5 ng/ml and 4.5 ng/ml, the measuring range (defined by the limit of detection and the maximum of the master curve) was 5–3000 pg/mL and the intra-assay coefficient of variation was < 8.6% relative SD. For assessment of testosterone, biotinylated monoclonal anti-testosterone antibodies (rabbit) were used at a concentration of 40 ng/mL, the measuring range (defined by the limit of detection and the maximum of the master curve) was 0.025-15 ng/mL, and the intra-assay coefficient of variation was < 5.7% relative SD. A radioimmunoassay was used to determine the concentrations of estradiol and testosterone.

### Adipocyte Size Analysis

The adipocyte size in the adipose tissue was measured according to previous studies (32). Briefly, adipocyte boundary segmentation was primarily conducted using AdipoCount software (33) and then was re-segmented manually to improve accuracy. Segmented adipocyte images were analyzed using ImageJ software (34). Image thresholds were adjusted from 55 to 255, and adipocyte sizes were analyzed with the “Analyze Particles” menu, with size=500- 30,000 µm^2^. All analyzed adipocyte size data were distributed into 1000 µm^2^ units.

### Insulin Tolerance Test and Oral Glucose Tolerance Test

For the insulin tolerance test (ITT), rats were fasted overnight and weighed. Blood sugar was measured using a glucometer before and 15, 30, 45, 60, 90 and 120 min after intraperitoneal insulin injection (0.5 IU/kg bodyweight; Humulin R, Eli Lilly and Company, USA). The time required for blood sugar levels to decline by 50% (t_1/2_) was calculated by linear regression and the K_ITT_ value was calculated by the following equation: K_ITT_ = 69.3/*t*_1/2_. For the oral glucose tolerance test (OGTT), rats were fasted overnight and weighed. Blood sugar was measured using a glucometer before and 30, 60, 90, and 120 min after intragastric glucose administration (2 g/kg body weight; 50% Dextrose Inj, Dai Han Pharm Co., Republic of Korea).

### Blood Pressure Measurement

Indirect measurement of systolic blood pressure (SBP) was carried out using the tail-cuff method (ML125; Powerlab, AD Instruments, Castle Hill, NSW, Australia). Before SBP measurement, the rats (24-week-old normal diet-fed rats and 15-week-old high-fat diet-fed rats) were placed in a warming chamber set at approximately 34 °C for 15 minutes, followed by placement in a plastic restrainer. To minimize stress-induced pressure changes, the animals were acclimated to the measurement conditions of the plethysmograph beforehand. SBP measurements were consistently taken between 10 AM and 12 PM. A minimum of five consecutive measurements were recorded using the data acquisition system (LabChart® software, version 7.1; ADInstruments, Colorado Springs, CO) to determine the mean SBP for each rat.

### Plasma Insulin and Free Fatty Acid Analysis

Blood was collected in EDTA tubes from rats fasted overnight, and plasma was separated using a centrifuge at 1500 x g for 10 min. Fasting insulin was assessed using a Rat Insulin ELISA Kit (RTEB0287, AssayGenie, Ireland) following the manufacturer’s instructions and measured using a SpectraMax ABS microplate reader (Molecular Devices, USA). The fasting free fatty acid level was assessed using a Free Fatty Acid Assay Kit (ab65341, Abcam, United Kingdom) following the manufacturer’s instructions and measured using a SpectraMax ABS microplate reader.

### ADSC Isolation and Adipogenic Differentiation

Perigonadal and inguinal adipose tissues were dissected from 14-to-15-week-old rats and minced using surgical blades. Minced tissues were digested in 10 mL of 0.05% collagenase type I (Gibco, USA)/HBSS (Gibco) for 30 min at 37 ℃ in a 5% CO_2_ incubator. Digested tissues were centrifuged at 2000 x g for 5 min, and the supernatant was discarded. Centrifuged stromal vascular fractions were filtered through 70 µm nylon mesh and transferred to new 1.5 mL tubes and red blood cells were lysed using eBioscience™ 1X RBC Lysis Buffer (Invitrogen). ADSCs were cultured in 20% fetal bovine serum (Gibco)/DMEM (Cytiva, USA) supplemented with 1% Pen/Strep (Gibco). ADSCs were grown to 90% confluence and differentiated into adipocytes using a StemPro™ Adipogenesis Differentiation Kit (Gibco) following the manufacturer’s instructions.

### Neutral Lipid-Positive Pixel Counts

Differentiated adipocytes were stained using BODIPY™ 493/503 (Invitrogen) and 9 images per each group were collected using an EVOS M7000 (Invitrogen) at the same locations. The color channels of the images were split, and green channel images were used for analysis. The threshold for measuring the ratio of green fluorescence positive pixels was set between 55 and 255, and only pixels within this range were counted.

### Statistical Analysis

Data were analyzed using Student’s t-test, which was performed using GraphPad Prism version 8.0.1 for Windows (GraphPad Software, USA, www.graphpad.com). When the p-value was lower than 0.05, the results were considered statistically significant. The exact meaning of the symbol is included in the figure legends.

## Results

### CYP17A1-Deficient Rats Showed Sex Abnormality and Obesity With Increased Subcutaneous Fat

Using the CRISPR-Cas9 system, the sequence following the *Cyp17a1* gene start codon was targeted and *Cyp17a1* KO rats were produced (S1 Fig a). PCR analysis and Sanger sequencing at the genomic DNA level were used to primarily validate the gene’s knockout and a 295-bp deletion at the target region was discovered (S1 Fig b–e). Males with CYP17A1 deficiency had a sex-reversed phenotype at their external genitalia due to CYP17A1’s involvement in steroidogenesis (Fig 1a–d), and the reproductive tracts of *Cyp17a1* KO rats had sexual abnormalities (Fig 1e–h). H&E staining confirmed that KO female rats had thinner uteri than wild-type female rats (224.4 µm vs. 903.5 µm; Fig 1i, j) and failed folliculogenesis and ovulation through the absence of mature follicles and corpus luteum (Fig 1k, l). Similarly, the testis of *Cyp17a1* KO male exhibited failed spermatogenesis (Fig 1m, n). Quantitative PCR analysis revealed reduced *Cyp17a1* mRNA expression in the *Cyp17a1* (+/-) group and completely abolished expression in the knockout (-/-) group (Fig 1o), further confirming the gene’s knockout. To assess the impact of *Cyp17a1* gene knockout on sex hormone production, ECLIA was used to evaluate the levels of testosterone and estradiol in the serum of wild-type and *Cyp17a1* KO rats. *Cyp17a1* knockout males and females were confirmed to have impaired testosterone and estradiol production, respectively (Fig 1p, q). As these *Cyp17a1* KO rats are sterile, we maintained the line by mating *Cyp17a1* (+/-) males and females and cryopreserved the embryos.

**Fig 1.**
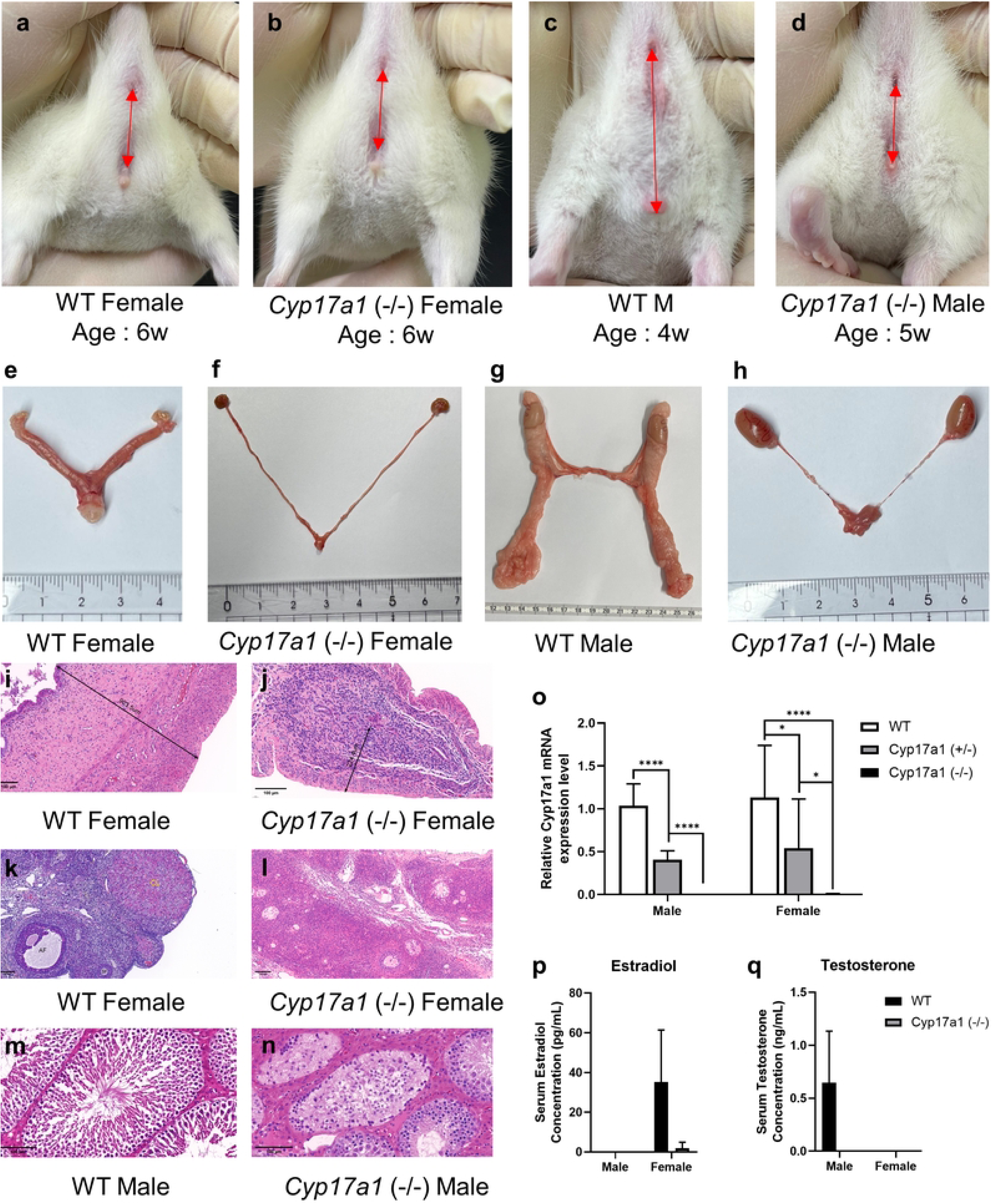
Phenotype of *Cyp17a1* knockout rats. **a-d)** External genitalia of rats. Red arrows indicate distance between anus and genitals. **e-h)** Reproductive tracts of rats. **i-j)** Representative images of H&E-stained uterus of female rats. Thickness of the uterus is indicated in the images. **k-l)** Representative images of H&E-stained ovary of female rats. SF: secondary follicle, AF: Antral follicle, CL: corpus luteum. **m-n)** Representative images of H&E-stained testis of male rats. Scale bars = 100µm. **o)** Cyp17a1 mRNA qPCR results of rats. Gonads from each group were used for qPCR analysis (n=3). The data were normalized to Actb and are presented as means±SEM. * p < 0.05, **** p < 0.0001, using Student’s t-test. p-q) LC-MS/MS result of serum estradiol and testosterone concentration in wild-type and Cyp17a1 knockout rats (n=3)

We also monitored the body weight of CYP17A1-deficient rats fed a high-fat diet since prior research had shown that mice with this gene defect have an obesity-related phenotype (30). *Cyp17a1* KO females had greater body weights than wild-type females (Fig 2a, c-d’, g) despite having equal muscle weights (Fig 2b). Interestingly, an increase in visceral and subcutaneous fat was observed in the *Cyp17a1* KO female group (Fig 2h, i) and an increased fat-to-body-weight ratio was also confirmed (Fig 2j, k). This trend was also observed in the male group, where, despite their body weight being comparable to that of the wild-type males (Fig 2a, e-f’, g), a similar or slightly greater amount of fat was accumulated in *Cyp17a1* KO males (Fig 2h, i). Interestingly, the ratio of subcutaneous fat to body weight rose dramatically in both the *Cyp17a1* KO male and female groups (Fig 2j, k). Furthermore, it was discovered that the ratio of subcutaneous fat to visceral fat was also considerably increased in both the *Cyp17a1* KO male and female groups (Fig 2l). In the group following a chow diet, this trend remained the same. Although body weight was not significantly different from that of wild-type females and was less than that of wild-type males (S2 Fig a, b), there was a statistically significant increase in the amount of fat that each *Cyp17a1* KO group had acquired, particularly in subcutaneous fat (S2 Fig c–l). In both the male and female *Cyp17a1* KO groups, hypertrophy of visceral and subcutaneous fat was observed by adipocyte size measurement using H&E staining (Fig 2m–p, S3 Fig a–d). Remarkably, these outcomes were shown to a similar extent in both the male and female groups that were given a chow diet (S3 Fig e–l).

**Fig 2.**
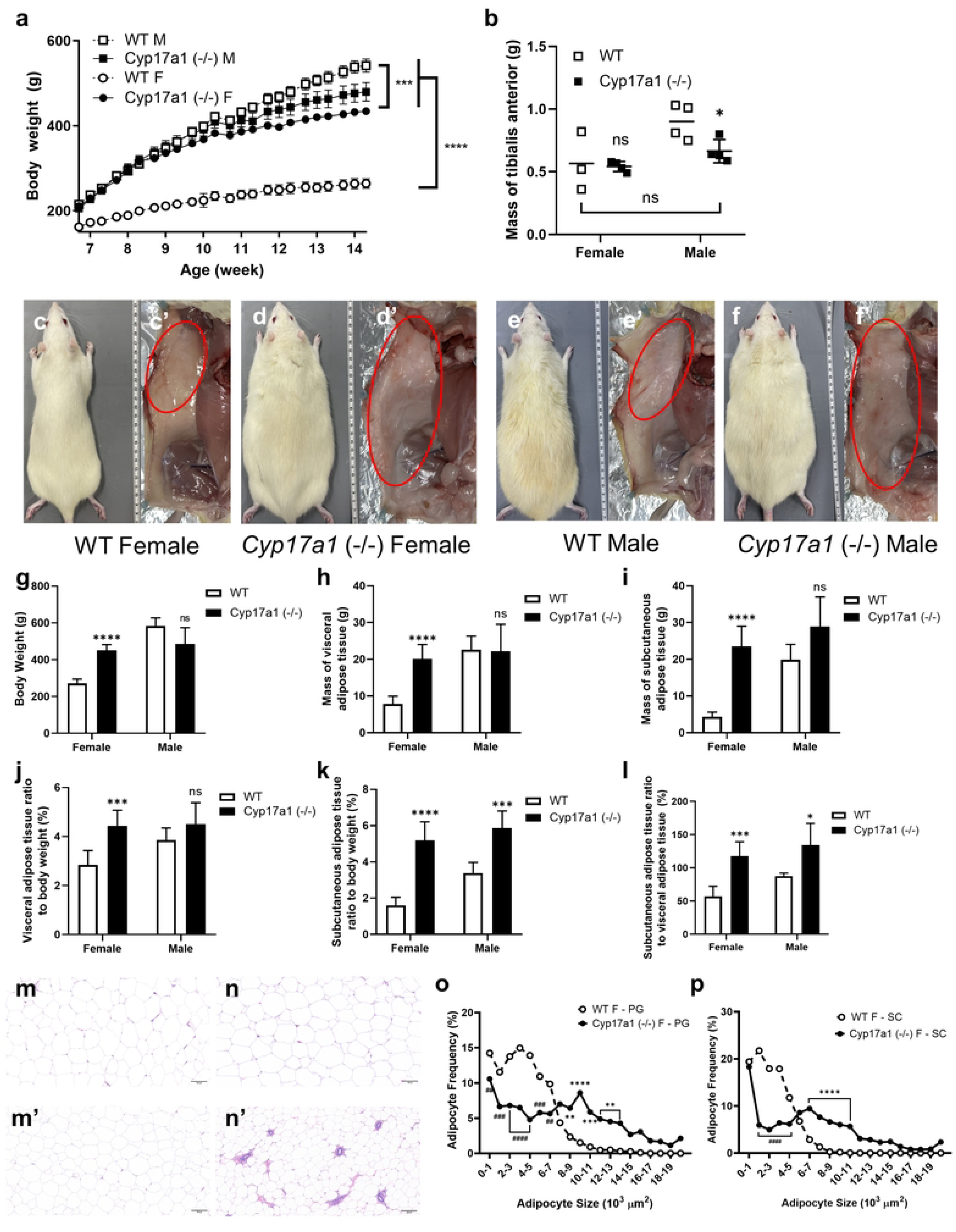
Fat distribution of *Cyp17a1* knockout HFD fed rats. **a)** Body weight measurement of HFD fed rats. Cyp17a1 knockout rats (n=8, male; n=12, female) and wild-type rats (n=4) were measured. ***p<0.001, ****p<0.0001, using Student t-test. **b)** Mass of skeletal muscle (tibialis anterior) of rats. (n=3, wild-type female; n=4, the other group) **c-f’)** Representative images of subcutaneous adipose tissue of HFD fed rats. Red circles indicate inguinal adipose tissue. **g-l)** Sampling results of rats. **(g)** Body weight, **(h)** mass of visceral adipose tissue, **(i)** subcutaneous adipose tissue, **(j, k)** relative mass of visceral and subcutaneous adipose tissue normalized to body weight. **(l)** Relative subcutaneous adipose tissue mass normalized to visceral adipose tissue. **m-n’)** Representative images of H&E-stained adipose tissue of HFD fed female rats. Visceral adipose tissue of Cyp17a1 knockout female rat **(m)**, wild-type female rat **(m’)**. Subcutaneous adipose tissue of Cyp17a1 knockout female rat **(n)** and wild-type female rat **(n’)**. Scale bar = 100µm. **o, p)** Adipocyte size analysis of perigonadal adipose tissue **(o)** and subcutaneous adipose tissue **(p)**. Images of H&E-stained adipose tissue were analyzed using ImageJ program (https://imagej.net/ij/). # p<0.05, ## p<0.01, ### p<0.001, #### p<0.0001, lower than wild-type. *p<0.05, **p<0.01, ***p<0.001, ****p<0.0001, higher than wild-type, using Student’s t-test, (n=12).

### CYP17A1 Deficiency in Rats Did Not Induce Metabolic Syndrome Despite Their Obesity and Hyperglycemia

As in previous results, *Cyp17a1* (-/-) rats showed signs of obesity, including a particular rise in subcutaneous fat and adipose tissue hypertrophy. Because CYP17A1 is involved in steroidogenesis and earlier studies have shown that changes in sex hormones are associated with insulin resistance (35–38), we conducted a series of assays to investigate the effect of CYP17A1 deficiency on metabolic syndrome. First, to examine if sexual maturation differs in metabolic syndrome between *Cyp17a1* KO rats and wild-type rats, we conducted an insulin tolerance test on chow-diet-fed rats at 5, 12, and 20 weeks old. At all three stages of age and in both sexes, there was no discernible difference in the *Cyp17a1* KO group’s and wild-type group’s responsiveness to insulin (S4 Fig a-i). When an insulin tolerance test was conducted on the high-fat-diet-fed group, males and females in the *Cyp17a1* KO group had significantly higher blood sugar levels than the wild-type group (Fig 3a, a’, d). However, the degree of drop in plasma glucose level after insulin injection was comparable to that in wild-type rats (Fig 3b, b’) and the K_ITT_ value, which indicates the percentage decline in plasma glucose concentration per minute, was also similar to that of wild-type (Fig 3c, c’).

**Fig 3.**
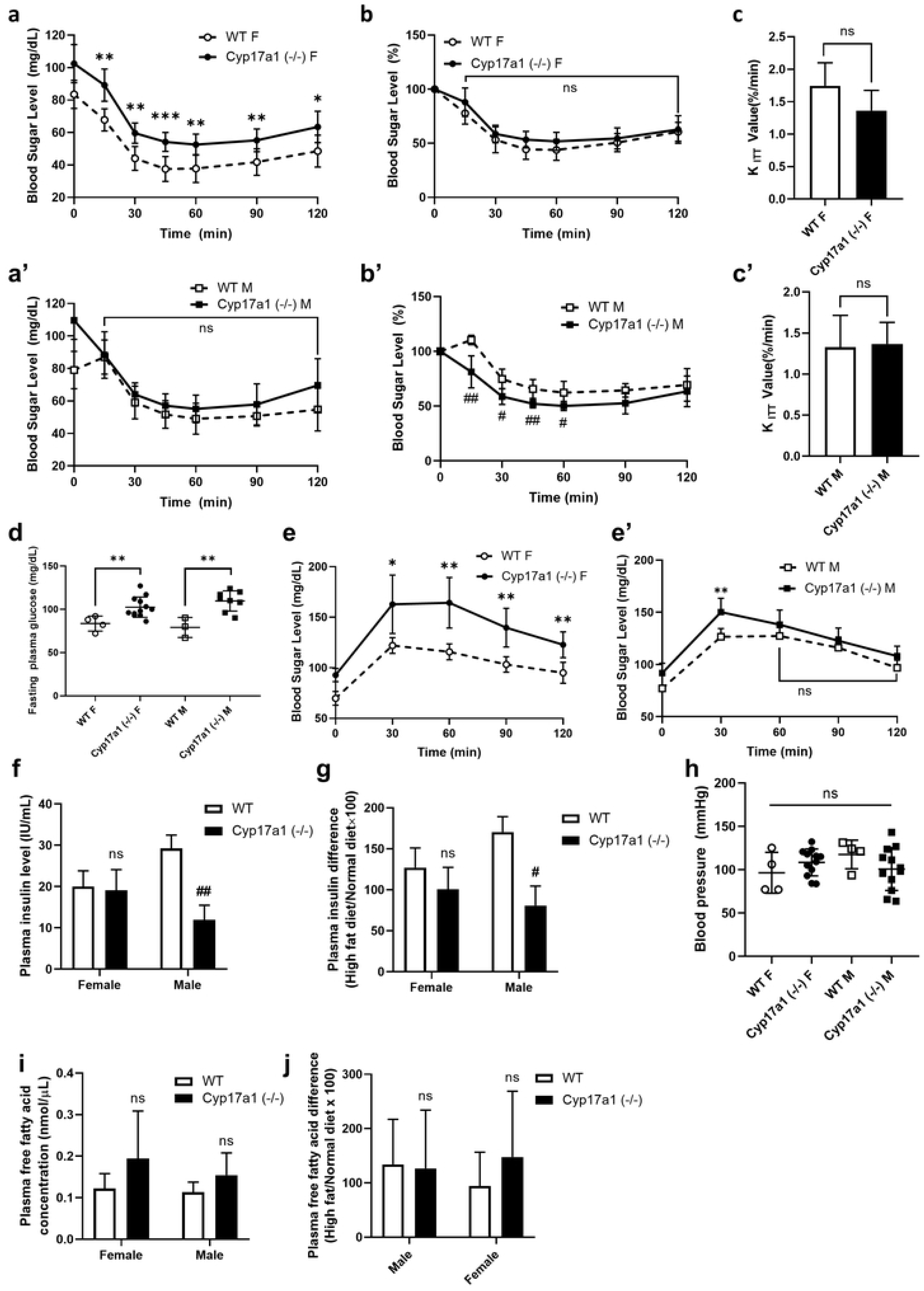
Metabolic analysis in *Cyp17a1* KO HFD fed female rats. **a-c’)** Results of insulin tolerance test (ITT) of HFD fed rats (n=12, *Cyp17a1* knockout group; n=4, wild-type group). **a, a’)** Blood glucose level after insulin injection and **b, b’)** percentage of blood glucose to fasting glucose level. ## p<0.01, lower than wild-type, *p<0.05, **p<0.01, ***p<0.001, higher than wild-type rats, using Student’s t-test. **c, c’)** K_ITT_ value which indicates percentage decline in blood glucose concentration per minute. **d)** Overnight-fast glucose level of HFD fed rats (n=12, *Cyp17a1* knockout group; n=4, wild-type group). **e, e’)** Results of oral glucose tolerance test (OGTT) of HFD fed rats. *p<0.05 **p<0.01, higher than wild-type, using Student’s t-test (n=12, *Cyp17a1* knockout group; n=4, wild-type group). **f)** Plasma insulin level of HFD fed rats analyzed with ELISA. **g)** Plasma insulin changes of HFD fed rats to chow diet-fed rats (n=6, *Cyp17a1* knockout group; n=3 wild-type group). **h)** Blood pressure of HFD fed rats. **i)** Plasma free fatty acid level of and **J)** free fatty acid level changes of HFD fed rats to chow diet-fed rats (n=6, *Cyp17a1* knockout group; n=3 wild-type group).

Subsequently, a glucose tolerance test was conducted on the rats to assess the insulin secretion response to glucose administration. In glucose tolerance tests conducted on *Cyp17a1* KO rats and wild-type rats fed a standard diet, no significant differences were observed between males and females in either group (S4 Fig j, k). Furthermore, males lacking the *Cyp17a1* gene displayed a reduced rise in blood glucose. However, when glucose tolerance tests were performed on rats fed a high-fat diet, significant differences were found between the female *Cyp17a1* KO group and the female wild-type group (Fig 3e). No such distinct differences were observed in the male *Cyp17a1* KO group, indicating a gender-specific response (Fig 3e’). When the plasma insulin concentration was measured, it was discovered that the insulin levels of the *Cyp17a1* KO females were similar to those of the wild-type females when they followed the chow diet and the high fat diet, while the insulin levels of the males did not increase when they followed the high fat diet, in contrast to the wild-type males (S4 Fig l, Fig 3f, g). Together with the earlier OGTT results, these data suggest that the reason *Cyp17a1* KO female rats’ blood sugar levels were higher was not due to an issue with insulin secretion, but rather due to hyperglycemia, which caused the blood sugar to be elevated overall.

To examine the metabolic syndrome caused by *Cyp17a1* KO in more detail, a blood biochemistry test was conducted. Although *Cyp17a1* KO females fed the chow diet showed higher triglyceride levels, they did not exceed the normal reference range (14.2 – 78.8 mg/dL, (39)) and the triglyceride, total cholesterol, HDL and LDL levels in the other groups were identical or even lower than those of wild-type females under both the high-fat diet and chow diet (Table 1, S2 Table). Both their free fatty acid level in plasma and blood pressure were the same in every group (Fig 3h–j, S4 Fig m). Considering previous research showed that elevated levels of free fatty acids lead to insulin resistance (10, 38), these findings provide more context for the observed hyperglycemia in *Cyp17a1* KO rats but equivalent insulin responses to wild-type rats. Furthermore, *Cyp17a1* KO rats did not exhibit the rise in plasma insulin concentration that is frequently seen in patients with metabolic syndrome, even when they were given a high-fat diet. Despite causing obesity-induced hyperglycemia, the *Cyp17a1* gene deletion did not affect insulin secretion or responsiveness. This shows that CYP17A1 deficiency cannot be considered to lead to a metabolic syndrome.

**Table 1.** Blood biochemistry of *Cyp17a1* knockout and wild-type rats with high fat diet.

| Blood Biochemistry Test (mg/dL) | Male |  |  | Female |  |  |
| --- | --- | --- | --- | --- | --- | --- |
|  | WT (n=3)<br>(mean±SEM) | *Cyp17a1* (-/-) (n=8)<br>(mean±SEM) | P-value (WT-<br>*Cyp17a1* KO) | WT (n=3)<br>(mean±SEM) | *Cyp17a1* (-/-) (n=12)<br>(mean±SEM) | P-value (WT-<br>*Cyp17a1* KO) |
| Triglyceride | 73±22.5 | 32±16.3 | 0.0047 | 57±11.8 | 36±17.9 | 0.0494 |
| Total Cholesterol | 74±7.8 | 74±20.1 | 0.9725 | 78±5.6 | 69±13.8 | 0.2177 |
| HDL | 20.8±1.31 | 22.4±3.87 | 0.4666 | 25.2±1.04 | 20.7±2.69 | 0.0063 |
| LDL | 10.9±1.23 | 6.5±1.71 | 0.0011 | 6.4±0.88 | 7.0±1.43 | 0.4344 |

### Enhanced Adipogenesis in ADSCs from *Cyp17a1* Knockout Rats

ADSCs were cultured from fat samples to investigate the etiology of obesity in the *Cyp17a1* KO group. Subsequently, adipogenic differentiation was induced and variations in adipocyte production were examined using neutral lipid staining. Based on the results of previous experiments which demonstrated more pronounced obesity and more prominent fat accumulation in the female group, ADSCs were isolated from visceral fat and subcutaneous fat in the high-fat-diet-fed female group and used in the following experiment about adipogenic potentials. Up until day 4, there was no discernible difference between the *Cyp17a1* KO group and the wild-type ADSCs isolated from visceral fat; however, on day 6, considerable differentiation was seen (Fig 4a, a’, c, c’, e). On the other hand, ADSCs isolated from subcutaneous fat in the *Cyp17a1* KO group consistently showed higher levels of neutral lipid formation, even before adipogenic stimulation (Fig 4b, b’, d, d’, f). To examine the adipogenic differentiation of ADSCs in further detail, RNA was isolated from differentiated ADSCs and qPCR was performed. During the 6-day period of induced adipogenic differentiation, explosive increases in the expression of early adipogenesis marker KLF5 and late adipogenesis markers PPARγ and C/EBPα were observed in both perigonadal and subcutaneous fat in the *Cyp17a1* KO group (Fig 4g–i’). Interestingly, while KLF5 expression was reduced in visceral-fat-derived ADSCs from *Cyp17a1* KO rats, it was enhanced in subcutaneous-fat-derived ADSCs from *Cyp17a1* KO rats.

**Fig 4.**
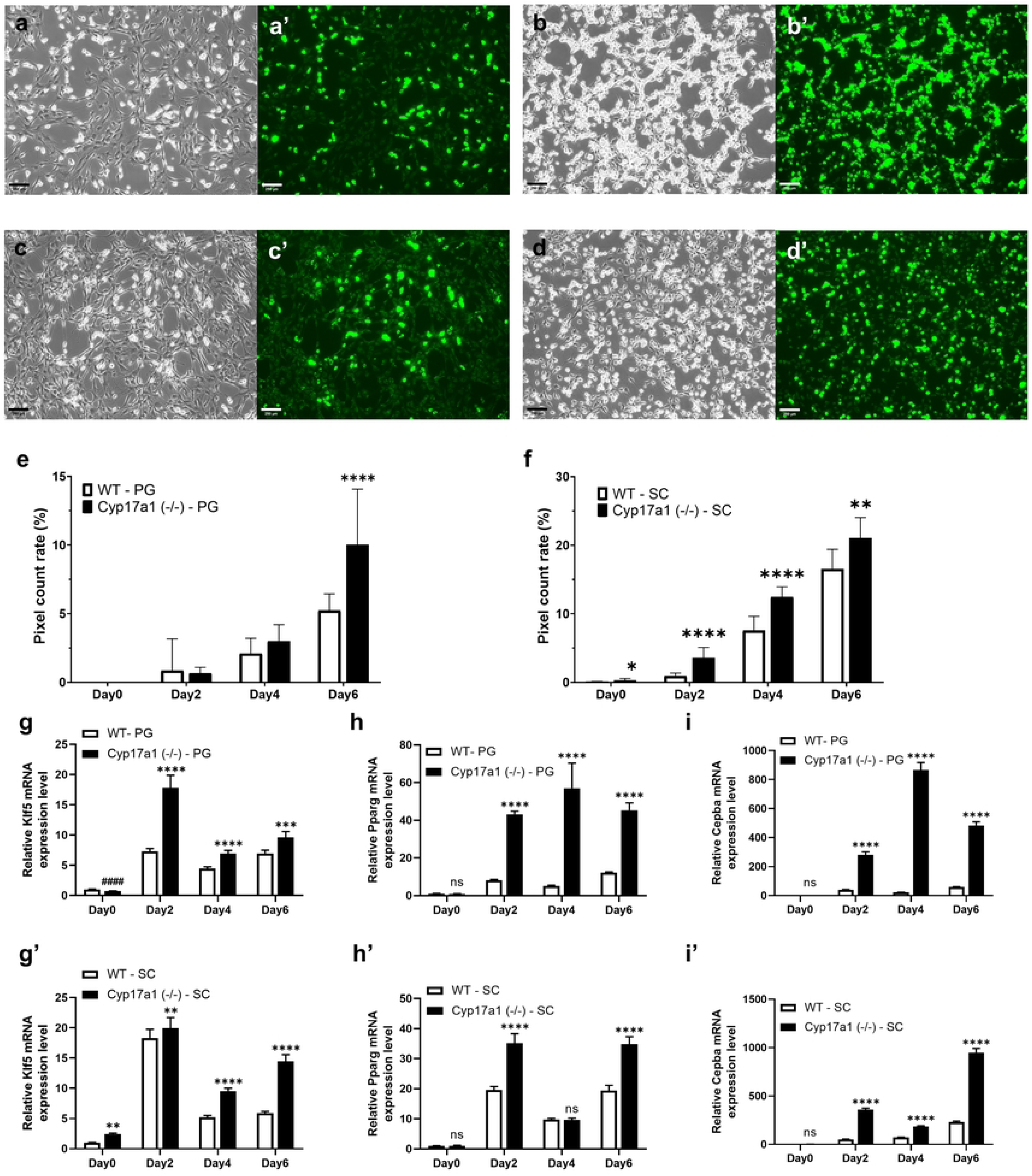
Adipogenesis using adipose-derived stem cell (ADSC) from *Cyp17a1* KO rats. **a-d’)** Representative images of adipogenesis using ADSCs from rats on day 6. Bright and fluorescent images of Cyp17a1 KO ADSCs from perigonadal adipose tissue **(a, a’)** and subcutaneous adipose tissue **(b, b’)**, and wild-type ADSC from perigonadal adipose tissue **(c, c’)** and subcutaneous adipose tissue **(d, d’)**. Scale bar=250µm. **e, f)** Results of neutral lipid positive pixel count using ImageJ program. Result of perigonadal adipose tissue derived stem cell **(e)** and subcutaneous adipose tissue derived stem cell **(f)**. *p<0.05, **p<0.01, ***p<0.001, ****p<0.0001 using Student’s t-test (n=9). **g-i’)** qPCR results of mRNAs from differentiated ADSCs on day 0, 2, 4, 6. Relative Klf5, Pparg, Cebpa mRNA expression level from differentiated ADSCs from perigonadal adipose tissue **(g-i)** and subcutaneous adipose tissue **(g’-i’)**. The data were normalized to Actb and are presented as means±SEM. #### p<0.0001, lower than wild-type, **p<0.01, ***p<0.001, ****p<0.0001, higher than wild-type, using Student’s t-test (n=6).

### Metabolic Shift from Visceral to Subcutaneous Fat Due to *Cyp17a1* Gene Knockout

RNA was then isolated from the previously sampled female adipose tissues and qPCR was conducted to confirm fat-related metabolism and inflammation. Remarkably, there was a confirmed decrease in the expression of several genes linked to fat metabolism in visceral fat and a rise in subcutaneous fat (Fig 5). When comparing the two lipogenesis-related genes *Srebf1* (SREBP-1c; sterol regulatory element-binding protein 1c) and *Dgat* (DGAT; diglyceride acyltransferase), the *Cyp17a1* KO group exhibited an overall higher gene expression level in subcutaneous fat but lower or no significant difference in visceral fat compared to wild-type group (Fig 5a, b). The *Cyp17a1* deletion group also exhibited higher expression levels in subcutaneous fat and lower or no significant differences in visceral fat when comparing the three lipolysis-related genes *Lipe* (HSL; hormone-sensitive lipase), *Prkaca* (PRKACA; protein kinase cAMP-activated catalytic subunit alpha), and *Pnpla2* (ATGL; adipose triglyceride lipase) (Fig 5c–e). In both visceral fat and subcutaneous fat, the expression levels of *Abhd5* (ABHD5; abhydrolase domain containing 5) and *G0s2* (G0S2; G0/G1 switch 2), a co-activator and regulator of ATGL respectively, were lower or did not differ significantly (Fig 5f, g), which means altered lipolysis activity in both adipose tissues is not regulated by the activity of other genes. The expression of three genes, *Mlxipl* (ChREBP; carbohydrate-responsive element-binding protein), *Slc2a4* (GLUT4; glucose transporter 4), and *Acaca* (ACACA; acetyl-CoA carboxylase alpha), related to glucose signaling lipogenesis, and two genes, *Irs1* (IRS1; insulin receptor substrate 1) and *Pik3ca* (PIK3CA; phosphatidylinositol-4,5-biphosphate 3-kinase, catalytic subunit alpha), related to the insulin signaling pathway, were examined to confirm whether this shift in fat metabolism to subcutaneous fat is related to glucose and insulin signaling. It was confirmed that visceral fat in *Cyp17a1* KO rats had decreased expression of genes related to glucose-induced lipogenesis, while subcutaneous fat showed an increase in this pathway (Fig 5h–j). Insulin signaling was found to be reduced in visceral fat of *Cyp17a1* KO rats and to be statistically comparable to that of wild-type rats in subcutaneous fat (Fig 5k, l), indicating reduced insulin-mediated lipogenesis in visceral fat and a normal insulin signaling pathway in subcutaneous fat. Subsequently, the expression levels of inflammation-associated genes *Tnf* (TNFα; tumor necrosis factor-alpha) and *Il6* (IL6, interleukin 6) were compared to examine inflammation in adipose tissue. In the subcutaneous fat of *Cyp17a1* KO rats, the expression of these genes was either the same or lower than in the wild-type rats, while in the visceral fat, *Il6* was more highly expressed and *Tnf* had lower expression levels (Fig 5m, n). Lastly, expression of the insulin resistance-related *Agt* (AGT; angiotensinogen) and *Adipoq* (AdipoQ; adiponectin) genes was found to be higher in subcutaneous fat and lower in visceral fat in *Cyp17a1* KO rats (Fig 5o, p).

**Fig 5.**
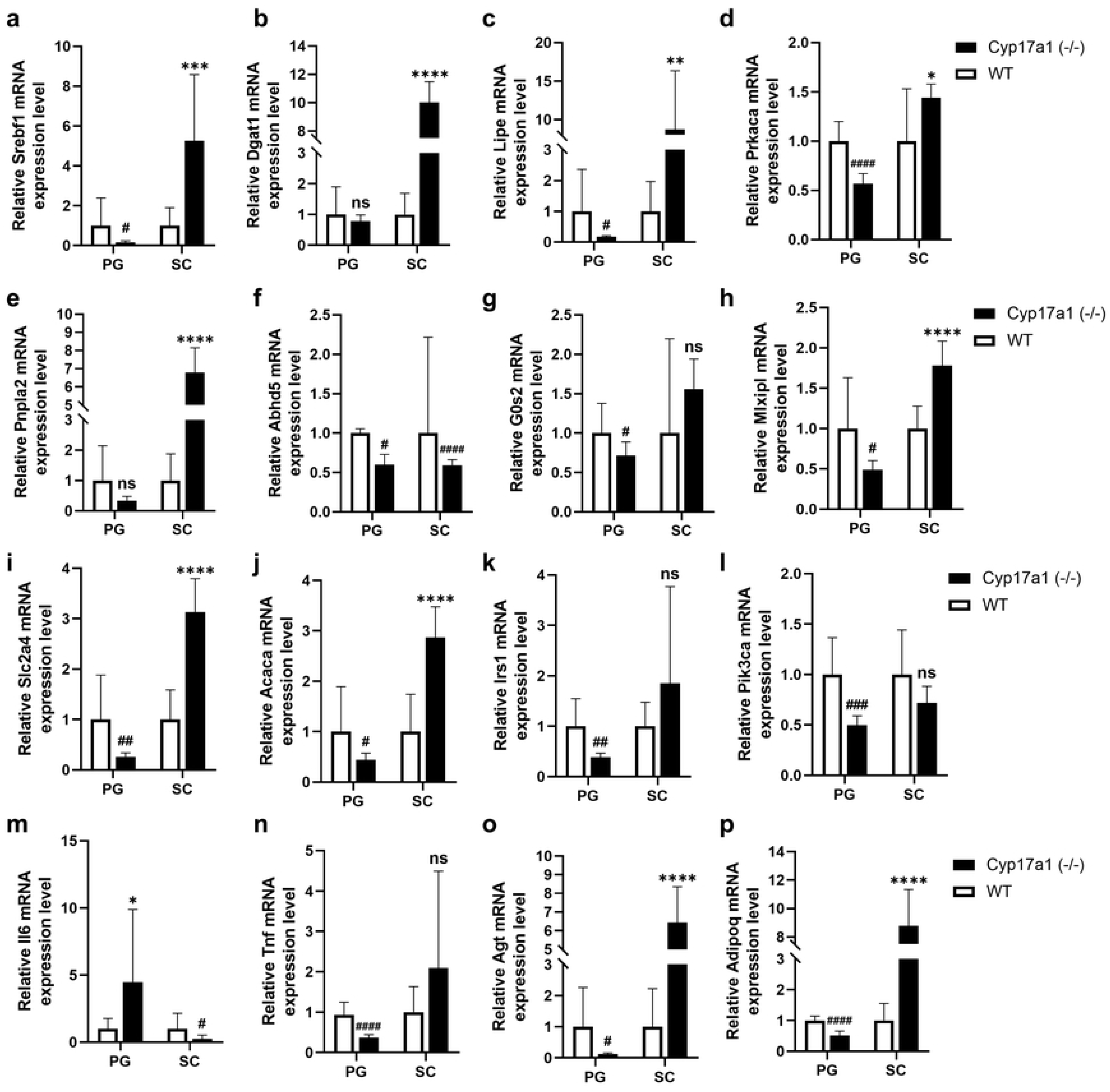
qPCR results of *CYP17A1* KO rats’ adipose tissue. **a, b)** Relative mRNA expression level of genes related to lipogenesis, Srebf1(SREBP-1c) and Dgat1 (DGAT). **c-e**) Relative mRNA expression level of genes related to lipolysis, Lipe (HSL), Prkaca (PRKACA), and Pnpla2 (ATGL). (f, g) Relative mRNA expression level of genes related to ATGL activity, Abhd5 (ABHD5) and G0s2 (G0S2). **h-j)** Relative mRNA expression level of genes related to glucose signaling mediated lipogenesis, Mlxipl (ChREBP), Slc2a4 (GLUT4) and Acaca (ACACA). **k, l)** Relative mRNA expression level of genes related to insulin-mediated glycerolipid metabolism, Irs1 (IRS1) and Pik3ca (PIK3CA). **m, n)** Relative mRNA expression level of genes related to adipose tissue inflammation, Il6 (IL6) and Tnf (TNFα). **o, p)** The data were normalized to Actb and are presented as means±SEM. Relative mRNA expression level of genes of adipokines, Agt(AGT) and Adipoq(AdipoQ). #p<0.05 ##p<0.01, ###p<0.001, #### p<0.0001, lower than wild-type, *p<0.05 **p<0.01, ***p<0.001, ****p<0.0001, higher than wild-type, using Student’s t-test (n=12).

## Discussion

This study explored the role of the *Cyp17a1* gene in obesity and its subsequent impact on metabolic diseases using a KO rat model, which was generated for the first time by CRISPR- Cas9. We conducted a detailed comparison of altered lipid and glucose metabolism between subcutaneous and visceral fat depots. We found that while *Cyp17a1* gene knockout induced obesity, metabolic syndrome did not appear, and that *Cyp17a1* gene knockout affected distinct metabolic shifts, underscoring its potential role in modulating disease progression in metabolic syndrome.

We observed an atypical rise in subcutaneous fat in our KO rat model resulting from the deletion of the *Cyp17a1* gene. Except for hyperglycemia, the *Cyp17a1* KO group did not show abnormalities in insulin response or production despite obesity and resulting adipocyte hypertrophy. Furthermore, there was no change in the concentration of free fatty acids in the blood, nor any difference when a high-fat diet was substituted for a normal diet. This suggests that deletion of the *Cyp17a1* gene may suppress insulin resistance, but further research involving a comprehensive analysis of glucose-lipid metabolism including skeletal muscle and liver (40–42) is necessary to confirm the effectiveness of the *Cyp17a1* KO model.

ADSCs were isolated and adipogenesis was stimulated to carry out a more thorough examination of obesity. Adipogenic gene expression and a greater differentiation rate were observed in ADSCs obtained from females lacking *Cyp17a1*. Although it is difficult to conclude from these data whether ADSCs directly develop into adipocytes in adipose tissues in vivo, the adipogenic potential is enhanced in the adipose tissue by *Cyp17a1* gene knockout in comparison to wild-type. Specifically, before triggering adipocyte differentiation, ADSCs from subcutaneous fat in the *Cyp17a1* KO group exhibited higher *Klf5* mRNA expression, whereas visceral fat showed lower expression. Considering KLF5 facilitates the induction of adipogenic gene expression by PPARγ and C/EBPα, the findings can be interpreted as in vitro data demonstrating that *Cyp17a1* knockout results in a shift in fat metabolism to subcutaneous depots (Fig 6).

**Figure 6.**
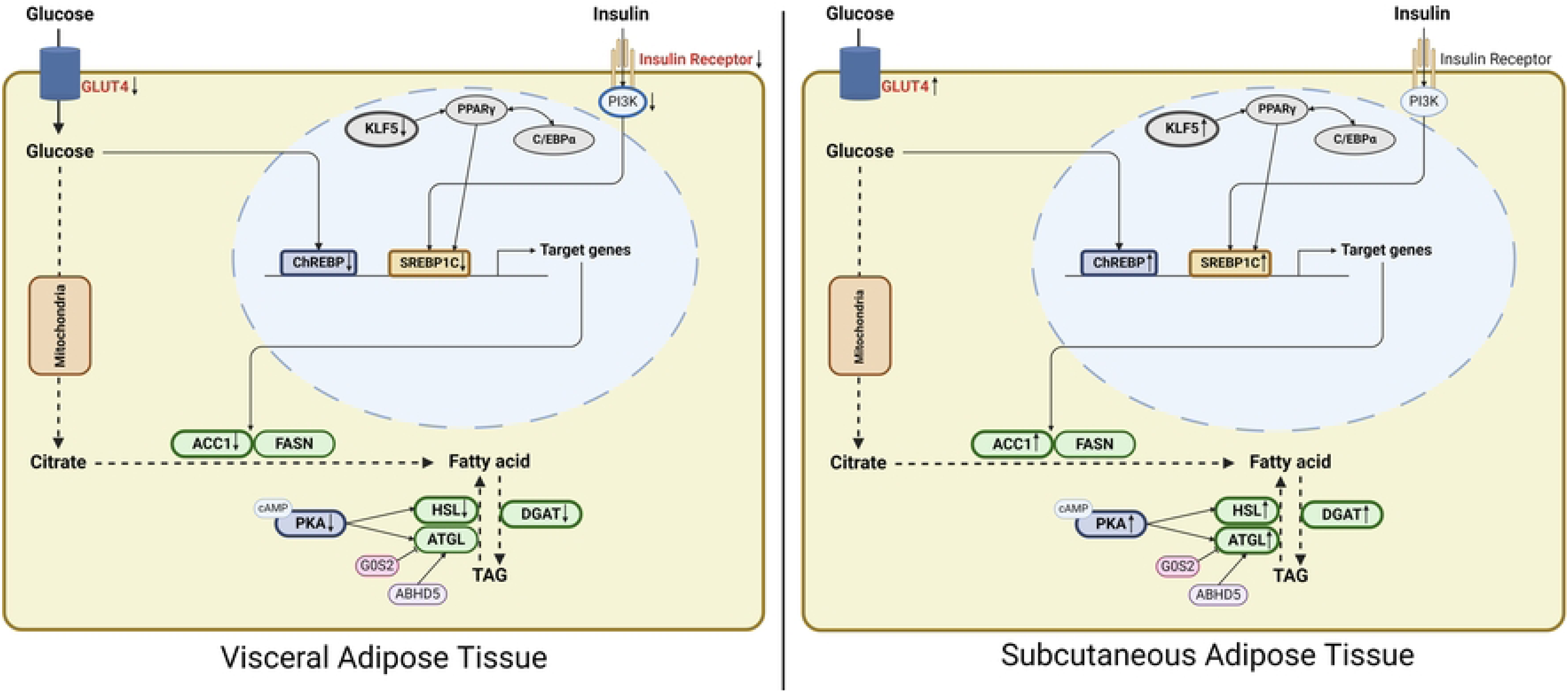
Graphical abstract of altered adipose tissue metabolism in *Cyp17a1* KO rats. Created in BioRender. Jang, G. (2024) BioRender.com/x87z478

Our findings indicate differential regulation of glucose and insulin signaling between visceral and subcutaneous adipose tissues, which may contribute to distinct metabolic functions. Specifically, glucose signaling was found to be reduced in visceral adipose tissue, whereas it was increased in subcutaneous adipose tissue. In contrast, insulin signaling was attenuated in visceral adipose tissue but remained relatively stable in subcutaneous adipose tissue. Interestingly, despite this stable insulin signaling in the subcutaneous tissue, we observed an upregulation in the expression of the downstream target SREBP1c. This suggests a potential redistribution of insulin signaling from visceral to subcutaneous fat depots. Furthermore, the observed shift in insulin signaling is likely linked to changes in *Klf5* gene expression, which is known to regulate lipogenesis in adipose tissue. The altered expression of *Klf5* may reflect an adaptive response in lipid metabolism, specifically in subcutaneous fat, and contribute to differential lipid storage and processing between these fat depots. Collectively, these findings suggest that the shifts in glucose and insulin signaling, along with changes in *Klf5* expression, are pivotal in driving the distinct lipid metabolic profiles observed in visceral and subcutaneous fat. This may provide a mechanistic basis for the differences in lipid metabolism and its associated metabolic outcomes between these two fat depots (Fig 6). As previous studies suggest that insulin resistance is caused by visceral fat dysfunction (15–17), it is assumed that the metabolic shift induced by *Cyp17a1* deletion protects against metabolic disorders despite obesity. Also, the findings that visceral fat secretes fewer genes linked to inflammation and insulin resistance, such as *Tnf* and angiotensinogen, and that subcutaneous fat tissue does not inflame but instead secretes a significant amount of adiponectin can also be interpreted as supporting the prior assumption.

Changes in steroid hormone levels may be involved in the healthy obesity trait observed in *Cyp17a1* KO rats. Numerous studies have demonstrated the connection between glucocorticoids and both insulin resistance and obesity (43–46). Excess glucocorticoids are believed to inhibit the immune system in adipose tissue by modulating macrophages and metabolism. Since the synthesis of glucocorticoids and sex hormones depends on the *Cyp17a1* gene involved in steroidogenesis, the disruption of hormone secretion in our *Cyp17a1* KO rats may contribute to their metabolically healthy obesity phenotype without insulin resistance. Consistent with our findings, another study demonstrated that steroidogenesis-related HSD11B1 deficiency prevents the development of metabolic syndrome, however, the study’s mechanism was not made explicit (47). Further investigation into the effects of glucocorticoids and other steroid hormones on metabolism in fat, skeletal muscle, the liver, etc. in *Cyp17a1* KO rats seems to be required in this respect. We cryopreserved the *Cyp17a1* KO embryos and hope our *Cyp17a1* KO line can be shared with other researchers and used in future studies.

In conclusion, in this study, we demonstrated that *Cyp17a1* deletion in rats via CRISPR- Cas9 caused obesity. Notably, the altered fat metabolism observed in the transition from visceral fat to subcutaneous fat suggests a protective mechanism. This metabolic shift may have played a key role in preventing the progression of a metabolic syndrome phenotype, despite obesity. While more investigation is needed to determine the impact of a prolonged high-fat diet on *Cyp17a1* KO rats and to elucidate the precise mechanism underlying the association between *Cyp17a1* deletion and metabolic syndrome, we anticipate that our research will further knowledge regarding the complex relationship between adipose tissue and metabolic syndrome.

## Supporting Information Caption

**S1 Fig.** Production of *Cyp17a1* knockout rats. **a)** Schematic image of CRISPR/Cas9 target site on rat *Cyp17a1* gene. Blue line indicates start codon site and red arrow indicates cut site. **b-d)** PCR analysis results of F0, F1 and F2. F0 and *Cyp17a1* (+/-) rats showed two bands, 877bp and 582bp, and wild-type rats and *Cyp17a1* (-/-) rat showed one band, 877bp upper band and 582bp lower band each. **e)** Sanger sequencing result of Cyp17a1 knockout F0 rat. Box indicates deleted sequence, 295bp. Blue letters, start codon.

**S2 Fig.** Sampling results of chow diet fed *Cyp17a1* knockout rats. **a)** Body weight measurement of chow diet fed rats age from 3 to 12 week (n=3). **b)** Body weight measurement of chow diet fed rats age from 12 to 20 week (n=6, *Cyp17a1* KO group; n=3, wild-type group). **c-f’)** Representative images of subcutaneous adipose tissue of chow deit fed rats. Red circles indicates inguinal (subcutaneous) adipose tissue. **g-l)** Sampling results of rats. **(g)** Body weight, **(h)** mass of visceral adipose tissue, **(i)** subcutaneous adipose tissue, **(j, k)** relative mass of visceral and subcutaneous adipose tissue normalized to body weight. **l)** Relative subcutaneous adipose tissue mass normalized to visceral adipose tissue (n=9, *Cyp17a1* KO male; n=8, *Cyp17a1* KO female, n=3; wild-type group). ns, not significant, *p<0.05, **p<0.01, ***p<0.001, ****p<0.0001, higher than wild-type, using Student’s t-test.

**S3 Fig.** Analysis of adipose tissue from Cyp17a1 knockout rats. **a, a’, b, b’)** Representative images of H&E stained perigonadal adipose tissue **(a, b)** and subcutaneous adipose tissue **(a’, b’)** of HFD fed male group. **c, d)** Adipocyte size analysis of perigonadal adipose tissue **(c)** and subcutaneous adipose tissue **(d)** from HFD fed male group. **e, e’, f, f’)** Representative images of H&E stained perigonadal adipose tissue **(e, f)** and subcutaneous adipose tissue **(e’, f’)** of chow diet fed female group. **g, h)** Adipocyte size analysis of perigonadal adipose tissue **(g)** and subcutaneous adipose tissue **(h)** from chow diet fed female group. **i, i’, j, j’)** Representative images of H&E stained perigonadal adipose tissue **(i, j)** and subcutaneous adipose tissue **(i’, j’)** of chow diet fed female group. **k, l)** Adipocyte size analysis of perigonadal adipose tissue **(k)** and subcutaneous adipose tissue **(l)** from chow diet fed female group. Scale bars=100µm. #p<0.05 ##p<0.01, ###p<0.001, #### p<0.0001, lower than wild- type, *p<0.05 **p<0.01, ***p<0.001, ****p<0.0001, higher than wild-type, using Student’s t-test (n=12).

**S4 Fig.** Metabolic analysis in chow diet fed *Cyp17a1* knockout rats. **a, d, g)** Results of insulin tolerance test (ITT) of chow diet fed female rats at 5 week age **(a)**, 12 week **(d)**, 20 week **(g)** (n=5, *Cyp17a1* knockout females at 5 week, n=6, *Cyp17a1* knockout females at the other age, n=3, wild-type females). **b, e, h)** Results of insulin tolerance test (ITT) of chow diet fed male rats at 5 week age **(b)**, 12 week (**e**), 20 week **(h)** (n=6, *Cyp17a1* knockout males, n=3, wild- type males). **c, f, i**) K_ITT_ value of chow diet fed rats at 5 week age (**c**), 12 week **(f)**, 20 week **(i)**. **j, k)** Results of oral glucose tolerance test (OGTT) of chow diet fed female **(j)** and male **(k)** rats. **l)** Plasma insulin level of chow diet fed rats analyzed with ELISA (n=6, *Cyp17a1* knockout group; n=3 wild-type group). **m)** Plasma free fatty acid level of chow diet fed rats (n=6, *Cyp17a1* knockout group; n=3 wild-type group).

